# Targeting distinct neuronal populations with AAV vectors reveals neurogenesis-associated anxiolytic effects of JNK inhibition

**DOI:** 10.64898/2026.10.30.763057

**Authors:** Anniina Jokela, Nikita Tiwari, Dani Flinkman, Soroush Abyari, Prasannakumar Deshpande, Jismi John, Eleanor Coffey

**Affiliations:** Turku Bioscience Centre, University of Turku and Åbo Akademi University, BioCity, Tykistökatu 6A, Turku-20520, Finland

**Author notes:** Corresponding author: Eleanor Coffey.

## Abstract

c-Jun N-terminal kinase (JNK) regulates stress responses and affective behaviour, but the neural populations mediating these effects remain unclear. Here, we delivered a nucleus-targeted JNK inhibitor to the mouse brain using blood-brain barrier-crossing adeno-associated virus vectors with distinct cellular tropisms. JNK inhibition produced strikingly different behavioural outcomes depending on the neuronal populations targeted. Only AAV-PHP.eB-hSyn1-mediated delivery reproduced the previously reported anxiolytic phenotype, whereas AAV.CAP-B10-hSyn1 increased locomotor activity and depressive-like behaviour and AAV.CAP-B10-mDlx reduced locomotion while enhancing object recognition memory. The anxiolytic response was accompanied by reduced hilar mossy cell abundance and an approximately two-fold increase in doublecortin-positive newborn granule neurons in the dentate gyrus. Notably, this phenotype mirrors our previous findings following selective JNK inhibition in adult hippocampal progenitor cells, implicating the neurogenic niche as a convergent substrate for anxiolytic responses to JNK inhibition. Proteomic analysis of plasma extracellular vesicles revealed widespread alterations in metabolic and inflammatory proteins upon JNK inhibition, despite neuron-restricted transgene expression. Together, these findings demonstrate that the behavioural and systemic consequences of JNK inhibition depend critically on the neural populations targeted and identify hippocampal neurogenesis-associated circuits involving mossy cells as candidate mediators of JNK-dependent anxiety regulation.

## INTRODUCTION

c-Jun N-terminal kinases (JNKs) are stress-activated protein kinases that belong to the mitogen-activated protein kinase (MAPK) family. JNK signaling controls various cellular functions such as apoptosis, differentiation, proliferation, and inflammation through phosphorylation of downstream targets which include, for example, transcription factors and mitochondrial proteins (Zeke et al., 2016). In the central nervous system (CNS), JNKs regulate brain morphogenesis, neuronal differentiation as well as axonal and dendritic remodeling during brain development, while also contributing to synaptic plasticity in the adult brain (Björkblom et al., 2005; Kuan et al., 1999; Li et al., 2007; Oliva et al., 2006; Tiwari et al., 2011). However, aberrant JNK activation has been implied in CNS disorders where it promotes, for example, neuroinflammation and dysregulated cell death, highlighting the roles of JNKs in both physiological and pathological processes (Corrales et al., 2021).

Moreover, JNKs have been implicated in neuropsychiatric disorders as, for example, *MKK7* upstream of JNK has been associated schizophrenia (Winchester et al., 2012) and JNK activity has been shown to regulate anxiety-like behaviours in several model organisms (Galeotti & Ghelardini, 2012; Hong et al., 2024; Zhou et al., 2023). We previously showed that knockout of *Jnk1*, intracerebral infusion with a JNK inhibitor, and retroviral delivery of a nuclear localised inhibitor of JNK (NLS-JBD) to the newborn neurons of the hippocampus all led to decreased anxiety-like behaviour in mice (Björkblom et al., 2005; Mohammad et al., 2018). Furthermore, in *Jnk1* knockout mice and with intracerebral JNK inhibitor infusion, adult hippocampal neurogenesis, a process associated with antidepressant drug action and anxiety-like behaviours (David et al., 2009; Revest et al., 2009), was increased (Mohammad et al., 2018), suggesting that JNK mediates its anxiolytic effects through the neurogenic niche.

Adeno-associated virus (AAV) vectors are non-pathogenic and non-integrating gene delivery tools used both in basic research and clinically approved gene therapies (Byrne et al., 2025; Samulski & Muzyczka, 2014). However, naturally occurring AAV serotypes poorly cross the highly restrictive blood-brain barrier (BBB) (Zincarelli et al., 2008), necessitating invasive administration methods in vector delivery to the brain. To overcome this limitation, various engineered AAV capsids have been developed with enhanced BBB permeability (Chan et al., 2017b; Goertsen et al., 2021). These vectors can be administered systemically through the blood stream, enabling minimally invasive brain-wide gene delivery to specific cell populations depending on the capsid tropism and the use of promoters. In addition to their versatility as research tools, these capsids also hold translational potential as non-invasive targeting of the human brain could facilitate gene therapy applications for brain disorders.

The main goal of this study is to establish whether using AAVs that deliver cargo across the BBB could be used to deliver a JNK inhibitor and mimic the anxiolytic effects elicited by gene knockout, intracranial infusion of inhibitor, and direct brain injection of retroviral vectors. By comparing different AAV capsids and cell-type-specific promoters, we wanted to identify the cell types and brain regions involved in this response.

## RESULTS

### Systemic delivery of JNK inhibitor using the AAV-PHP.eB capsid and the synapsin 1 promoter reduces anxiety like behaviour in mice

To address whether systemic delivery of a JNK inhibitor could replicate the anxiolytic effects of direct brain injection, we first utilized the AAV-PHP.eB vector combined with the synapsin 1 promoter (Chan et al., 2017b) (Fig. 1a). C57BL/6 mice underwent retro-orbital injection with 1×10^10^ vg of AAV-PHP.eB-hSyn1-mRuby2 or AAV-PHP.eB-hSyn1-mRuby2-JBD (1-277) (Fig. 1a).

**Figure 1.**
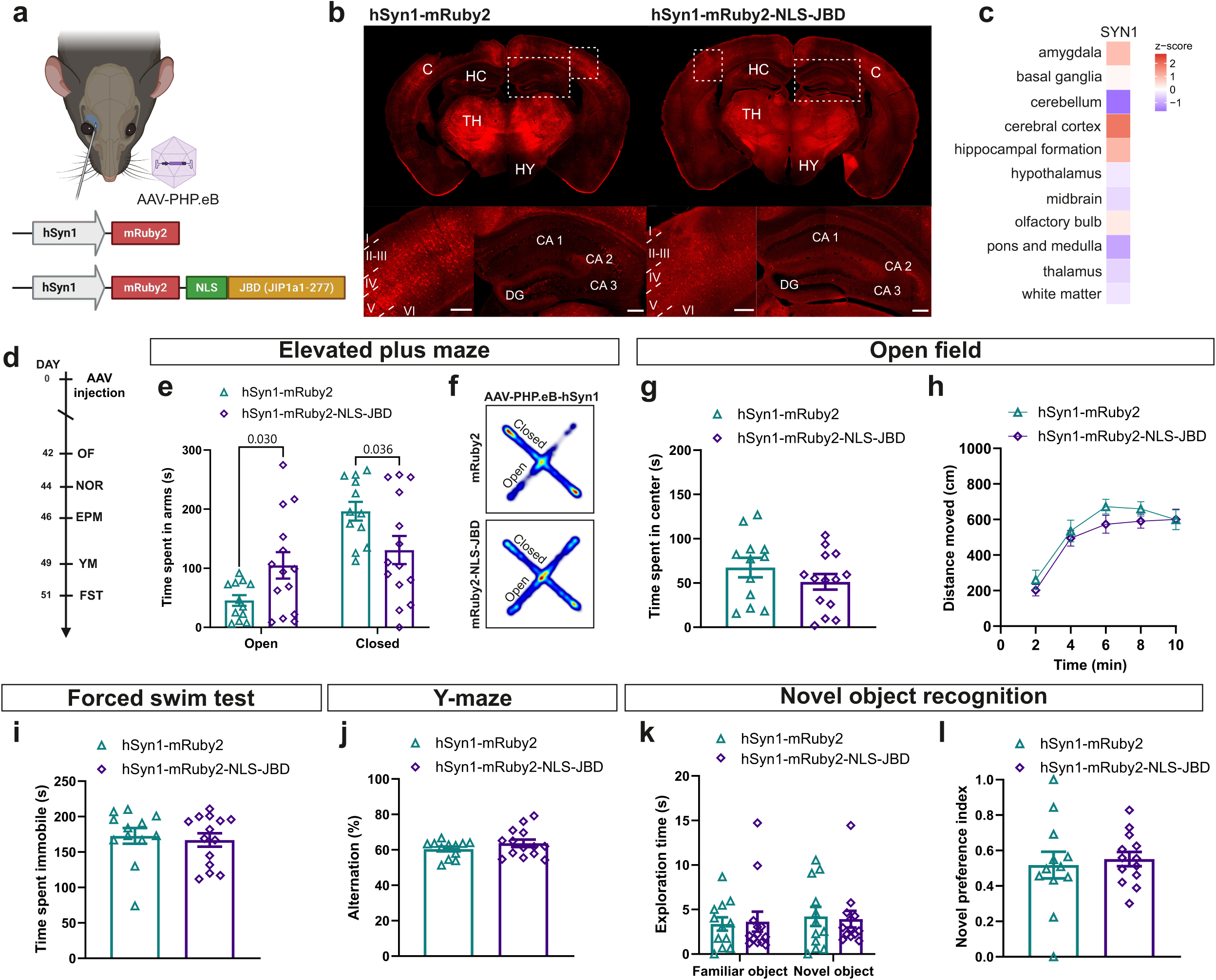
Brain-wide neuronal expression of a genetically encoded JNK inhibitor following systemic AAV-PHP.eB vector administration reduces anxiety-related behaviour in mice. **a)** 1×10^10^ vg of AAV-PHP.eB vectors were intravenously delivered into mice with the retro-orbital injection. Vectors used consisted of the hSyn1 promoter with either only the fluorescent reporter mRuby2 (control vector hSyn1-mRuby2) or mRuby2 with the nuclear JNK inhibitor sequence NLS-JBD (inhibitor vector hSyn1-mRuby2-NLS-JBD). **b)** Brain tissue expression of the synapsin-1 protein in the mouse brain from the Human Protein Atlas shows cerebral cortex, hippocampus, and amygdala as regions with the highest expression. **c)** Representative fluorescence images of mRuby2 from hSyn1-mRuby2 and hSyn1-mRuby2-NLS-JBD transduced mice based on the mRuby2 signal in the mouse brain eight weeks after AAV administration. Cortex and hippocampus are highlighted in the close-up images with cortical layers 1-5 and hippocampal regions labelled. **d)** Timeline of behavioural testing used in the experiment. **e-f)** Mice injected with the inhibitor vector hSyn1-mRuby2-NLS-JBD spent more time in the open arms and less time in the closed arms of the elevated plus maze compared to mice injected with the control vector hSyn1-mRuby2, indicating a reduction in anxiety-related behaviour. **(g)** In the open field test, there were no significant differences between the groups in time spent in the center of the arena or **h)** in total distance travelled, latter of which shows the locomotion of the mice was unaffected. **i)** Time spent immobile in the forced swim test shows no changes in depressive-like behaviour. **j-l)** Expression of JNK inhibitor did not affect the memory of the mice as evident in the unchanged spontaneous alternation in the Y-maze (j)) and in the time mice spent exploring objects **k)** and the novel preference index **l)** in the novel object recognition test. Data show mean +/-SEM for n=12 PHP.eB-hSyn1-mRuby2 mice and n=14 PHP.eB-hSyn1-mRuby2-NLS-JBD mice. Statistical analysis was done with Student’s two-tailed t-test or with two-way ANOVA. Abbreviations: CA=Cornu Ammonis, C=cortex, DG=dentate gyrus, HC=hippocampus, HY=hypothalamus, TH=thalamus, EPM=elevated plus maze, FST=forced swim test, NOR=novel object recognition test, OF=open field, YM=Y-maze.

Eight weeks later, they underwent behaviour testing (Fig. 1d). mRuby2 transgene expression was visible in the cortex, hippocampus and thalamus (Fig. 1b) Consistent with the known tropism of PHP.eB and synapsin 1 expression (Finneran et al., 2021; Kügler et al., 2003), mRuby2 expression for both vectors (hSyn1-mRuby2 and hSyn1-mRuby2-NLS-JBD), was observed throughout the cortical layers (Fig. 1b). In the hippocampus, mRuby2 was expressed in the dentate gyrus hilar cells and the granule cell layer. mRuby2 was also found in the pyramidal cell layer of CA1, CA2, and CA3 with strongest expression in the CA2 region as previously reported with AAV-PHP.eB (Mathiesen et al., 2020). This distribution corresponds with the expression of *synapsin 1* mRNA in mouse brain (Fig. 1c) (Sjöstedt et al., 2020a). Notably, the level of expression was lower for the mRuby2-NLS-JBD vector compared to the vector lacking NLS-JBD.

We next measured the effect of hSyn1-mRuby2 or hSyn1-mRuby2-NLSJBD on behaviour. Mice underwent a test battery that included the elevated plus maze, and forced swim test to measure affective behaviours, the open field to monitor locomotion and the Y-maze and novel object recognition tests to measure short-term spatial working memory and recognition memory. In the elevated plus maze, mRuby2-NLS-JBD expression induced a significant increase (> 2-fold) in the time spent in the open arms, indicating that these mice exhibited lower anxiety-like behaviour (Fig 1e-f). However, the JNK inhibitor (PHP.eB-hSyn1-mRuby2-NLS-JBD) did not significantly alter mouse behaviour in any of the other tests (Fig. 1g-l). These results indicate that inhibition of nuclear JNK in neurons that are targeted by the PHP.eB capsid, reduces anxiety-like behaviour without affecting locomotion or basic short-term memory tasks in mice.

### Systemic delivery of JNK inhibitor using the AAV.CAP-B10 capsid and the synapsin 1 promoter increases locomotion and depressive-like behaviour in mice

We next repeated the experiment using the AAV.CAP-B10 capsid. This capsid reportedly shows enhanced neuronal tropism and reduced transduction of liver cells compared to the AAV-PHP.eB (Goertsen et al., 2021). As previously, the vectors were delivered retro-orbitally into the bloodstream of mice (Fig. 2a). Eight weeks after injection of 3×10^11^ vg of vector, broad expression was observed in the cortex, thalamus and hippocampus (Fig. 2b). Compared to the AAV-PHP.eB-hSyn1 vector, cortical expression of AAV.CAP-B10-hSyn1-mRuby2 was more prominent in layer 5 pyramidal neurons. Otherwise the expression of both vectors was similar, and AAV.CAP-B10-hSyn1 was also expressed in the hilus and granule cell layer of the dentate gyrus, as well as the hippocampal pyramidal layer in CA1, CA2 and CA3.

**Figure 2.**
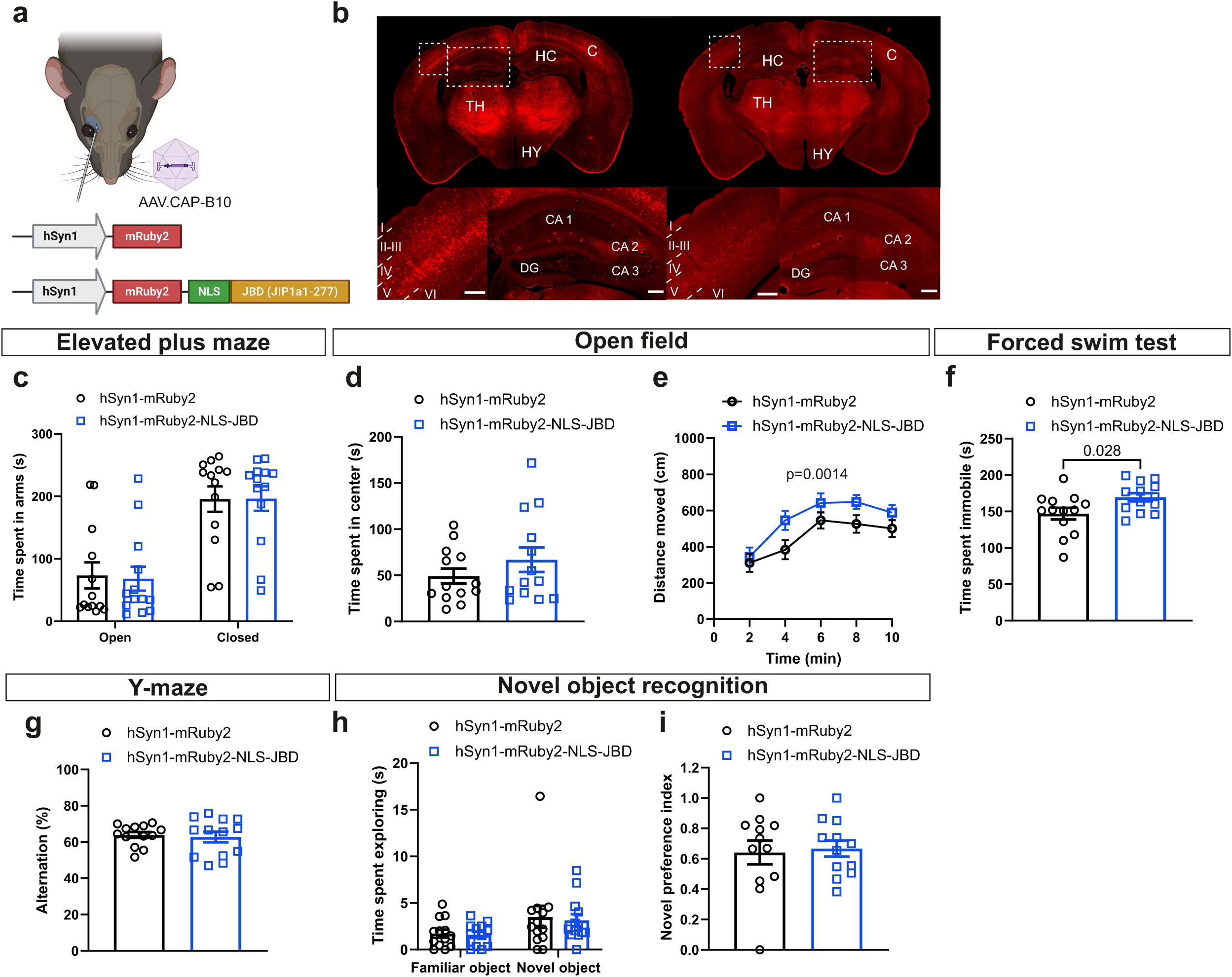
Brain-wide neuronal expression of a genetically encoded JNK inhibitor following systemic AAV.CAP-B10 vector administration leads to an increase in depressive-like behaviour in mice. **a)** 3×10^11^ vg of AAV.CAP-B10 vectors expressing either only mRuby2 (control vector hSyn1-mRuby2) or mRuby2 with the nuclear JNK inhibitor sequence NLS-JBD (inhibitor vector hSyn1-mRuby2-NLS-JBD) under the hSyn1 promoter were intravenously delivered into mice with the retro-orbital injection **b)** Representative images of hSyn1-mRuby2 and hSyn1-mRuby2-NLS-JBD vector expression based on the mRuby2 signal in the mouse brain eight weeks after AAV administration. Cortex and hippocampus are highlighted in the close-up images with cortical layers 1-5 and hippocampal regions labelled. **c)** In the behaviour testing of hSyn1-mRuby2 and hSyn1-mRuby2-NLS-JBD mice, expression of the nuclear JNK inhibitor did not affect anxiety-related behaviour of the mice as seen in the time spent in the arms of the elevated plus maze test and **d)** in time spent in the center of the open field test. **e)** Locomotion of the mice was also unaffected as seen in the total distance travelled in the open field test. **f)** In the forced swim test, hSyn1-mRuby2-NLS-JBD mice spent more time immobile in the water, indicating increased depressive-like behaviour. **g)** Memory and cognition of the mice was unaffected based on the spontaneous alternation in the Y-maze test and **h-i)** time spent exploring the objects and the novel preference index in the novel object recognition test. Data show mean +/-SEM for n=13 CAP-B10-hSyn1-mRuby2 mice and n=13 CAP-B10-hSyn1-mRuby2-NLS-JBD mice. Statistical analysis was done with Student’s two-tailed t-test or with two-way ANOVA. Abbreviations: CA=Cornu Ammonis, C=cortex, DG=dentate gyrus, HC=hippocampus, HY=hypothalamus, TH=thalamus.

Interestingly, mRuby2-NLS-JBD delivered using the CAP-B10 capsid did not alter anxiety-like behaviour (Fig 2c-d), in contrast to the results the PHP.eB capsid (Fig. 1e). Rather, with the CAP-B10-hSyn1-mRuby2-NLS-JBD vector, mice displayed increased locomotion in the open field compared to control mice expressing mRuby2 alone (Fig. 2e). Similarly, in the forced swim test, these mice spent more time immobile compared to control mice (Fig. 2f), suggesting a depressed-like state. In the Y-maze and novel object recognition tests, CAP-B10-hSyn1-mRuby2-NLS-JBD expression did not alter the behaviour indicating that there was no effect on short term spatial working memory or episodic memory of the mice (Fig. 2g-i). These results suggest that inhibiting nuclear JNK in the brain cells targeted by the AAV.CAP-B10 capsid increases locomotion and depressive-like behaviour. This distinguishes it from the results obtained with the AAV-PHP.eB capsid (Fig. 1), where there was a clear anxiolytic response.

### Selective targeting of JNK inhibitor to inhibitory neurons decreases locomotion

Since we saw a reduction in anxiety-like behaviour with the PHP.eB capsid but not with CAP-B10, though in both cases expression was in neurons (i.e. under the hSyn1 promoter), we wanted to test what would happen if we targeted inhibitory neurons. Therefore, we used the mDlx promoter. Mice were transduced retro-orbitally with 3×10^11^ vg of the AAV.CAP-B10-mDlx-NLS-mRuby2 vectors (Fig. 3a). After 10 weeks of expression, a weak signal was obtained for mRuby2 in the hippocampus and cortex (Fig. 3b). The low signal was consistent with DLX5/DLX6 mRNA expression in these regions in the mouse brain (Fig. 3c). More prominent signal was present in the anterior olfactory nucleus and the olfactory bulb (Fig. 3b). In the cortex, mRuby2-positive cells were most abundant in the layers 2/3 and 4 and sparser in the deeper layers, whereas in the hippocampus, cells expressing these vectors were mainly located in the hilus of dentate gyrus and in the pyramidal layer of CA3. Strong expression in the olfactory bulb was expected based on the RNA expression of DLX5 and DLX6 (Fig. 3c) (Sjöstedt et al., 2020b).

**Figure 3.**
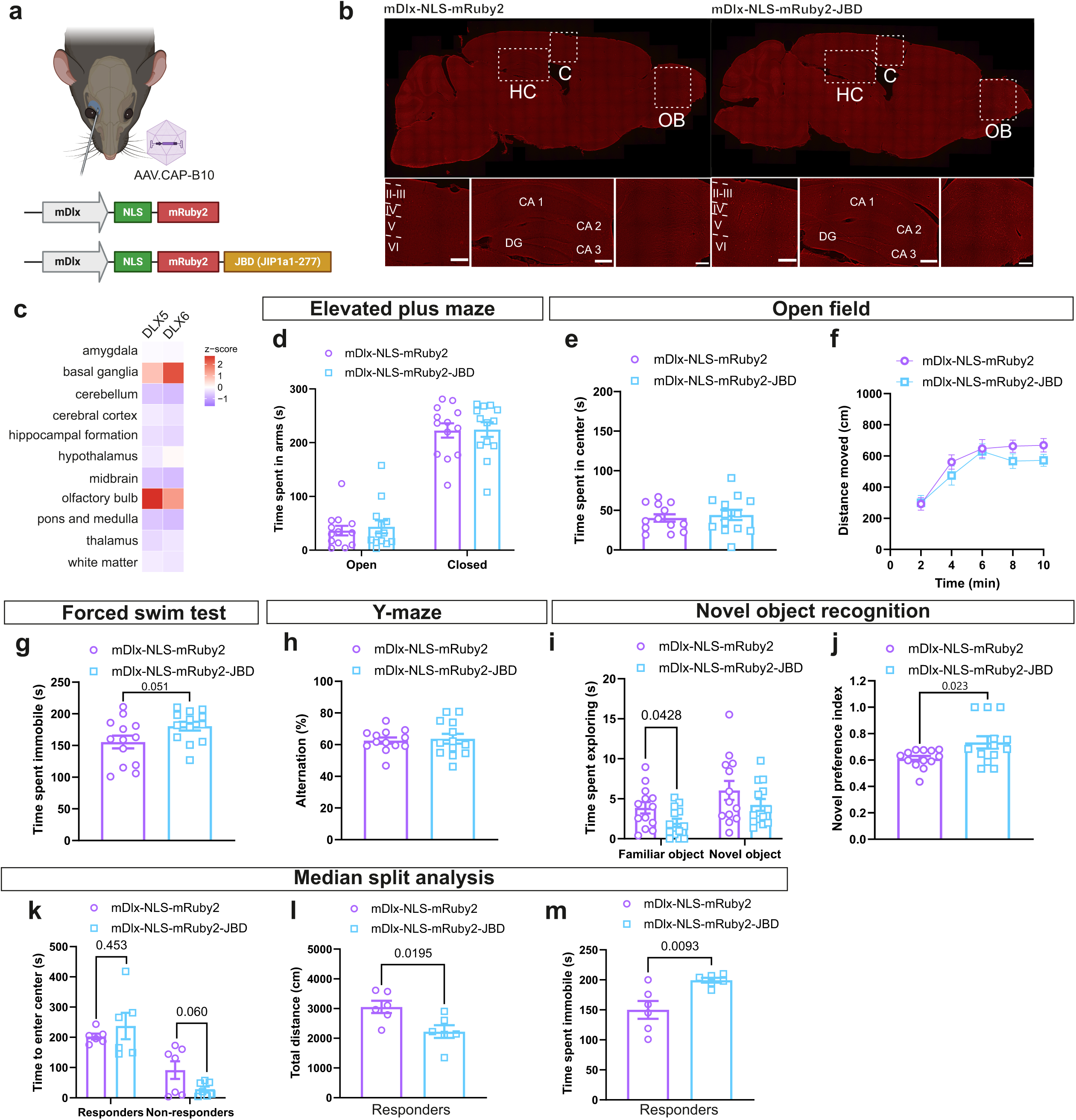
Expression of a genetically encoded JNK inhibitor in the inhibitory neurons of the mouse brain following systemic AAV.CAP-B10 vector administration improves recognition memory but impairs locomotion and increases depressive-like behaviour. **a)** 3×10^11^ vg of AAV.CAP-B10 vectors expressing either only NLS-mRuby2 (control vector mDlx-NLS-mRuby2) or mRuby2 with the nuclear JNK inhibitor sequence NLS-JBD (inhibitor vector mDlx-NLS-mRuby2-JBD) under the mDlx promoter were intravenously delivered into mice with the retro-orbital injection. **b)** Representative images of mDlx-NLS-mRuby2 and mDlx-NLS-mRuby2-JBD vector expression based on the mRuby2 signal in the mouse brain 10 weeks after AAV administration. In addition to cortex and hippocampus, olfactory bulb is highlighted in the close-up images as **c)** based on Human Protein Atlas data, it has a high expression of DLX5/6 in the mouse brain. In the behaviour testing of mDlx-NLS-mRuby2 and mDlx-NLS-mRuby2-JBD mice, **d)** time spent in the arms of the elevated plus maze test and **e)** time spent in the center of the open field test were unaffected. Additionally, **f)** there was no difference between the groups in distance moved in the open field test. **g)** In the forced swim test, there was a non-significant trend of mDlx-NLS-mRuby2-JBD spending more time immobile in the water. **h)** Spatial working memory was unaffected in the mice as seen in the Y-maze spontaneous alternation, however, **i-j)** expression of the JNK inhibitor seemed to improve recognition memory as mDlx-NLS-mRuby2-JBD mice had a higher novel preference index compared to control vector mice. **k)** After the control and inhibitor mouse groups were divided into responders and non-responders in a median split analysis based on time the mice took to enter the center of the open field arena, we found that in mice that took longer to enter the center, **l)** mDlx-NLS-mRuby2-JBD mice moved less in the open field and **m)** spent more time immobile in the forced swim test compared to the mDlx-NLS-mRuby2 mice, indicating that expression of a JNK inhibitor in the inhibitory neurons impairs locomotion and increases depressive-like behaviour. Data show mean +/-SEM for n=13 CAP-B10-mDlx-NLS-mRuby2 mice and n=13 CAP-B10-mDlx-NLS-mRuby2-JBD mice. Statistical analysis was done with Student’s two-tailed t-test or with two-way ANOVA. Abbreviations: CA=Cornu Ammonis, C=cortex, DG=dentate gyrus, HC=hippocampus, OB=olfactory bulb.

Six weeks after AAV.CAP-B10-mDlx vector administration, mice expressing mRuby2-NLS-JBD showed no changes in anxiety-like behaviours (Fig. 3d-e). At this stage (6 weeks), there was no locomotor difference (Fig. 3f), though by 10 weeks NLS-mRuby2-JBD-expressing mice did show significantly reduced locomotion (p=0.012; Supplementary Fig. 1a). In the forced swim test, there was a trend towards increased time immobile (Fig. 3g), which would reflect behavioural despair, however this was not significant (p=0.051). During the memory tests, mice performed equally well to control mice in the Y-maze, however, in the novel object recognition test the mDlx-NLS-mRuby2-JBD mice displayed a higher novel preference index (Fig. 3h-j), without any difference in distance moved (Supplementary Fig. 1b). Together these data suggest that nuclear inhibition of JNK in inhibitory neurons may improve recognition memory, though upon longer expression times e.g. 10 weeks, it may affect locomotion.

In order to uncover phenotypes that may have been masked by biological heterogeneity, the AAV.CAP-B10-mDlx-NLS-mRuby2 groups were divided into responders and non-responders based on the time the mice took to enter the center in the open field test (Fig. 3k). This revealed that mRuby2-NLS-JBD expressing mice moved significantly less in the open field (Fig. 3l) and spent a significantly longer time immobile in the forced swim test (Fig. 3m) six weeks after AAV injection. These results further indicate that inhibition of nuclear JNK in the inhibitory neurons in the mouse brain leads to a decrease in locomotion and induces depressive-like behaviour in susceptible mice.

### AAV-PHP.eB-hSyn1-mRuby2-NLS-JBD decreases hilar mossy cell number and increases DCX+ granule cell number in dentate gyrus

From among the tested vectors with different cellular tropisms (Fig. 1-3), only PHP.eB-hSyn1-mRuby2-JBD induced an anxiolytic response, replicating our earlier findings upon infusion of the JNK inhibitor peptide (Mohammad). We therefore evaluated whether this vector affected adult hippocampal neurogenesis. For this, we examined if PHP.eB-hSyn1-mRuby2-JBD expression altered expression of neurogenic cell markers in the hilus and dentate gyrus of the ventral hippocampus. First, we analysed the neural progenitor cell population by counting the proportion of transduced cells that were positive for the mini-chromosome maintenance-2 (MCM2) marker (Fig. 4a). There was no significant difference in the % of transduced cells that were MCM2+ when mRuby2-NLS-JBD was expressed.

**Figure 4.**
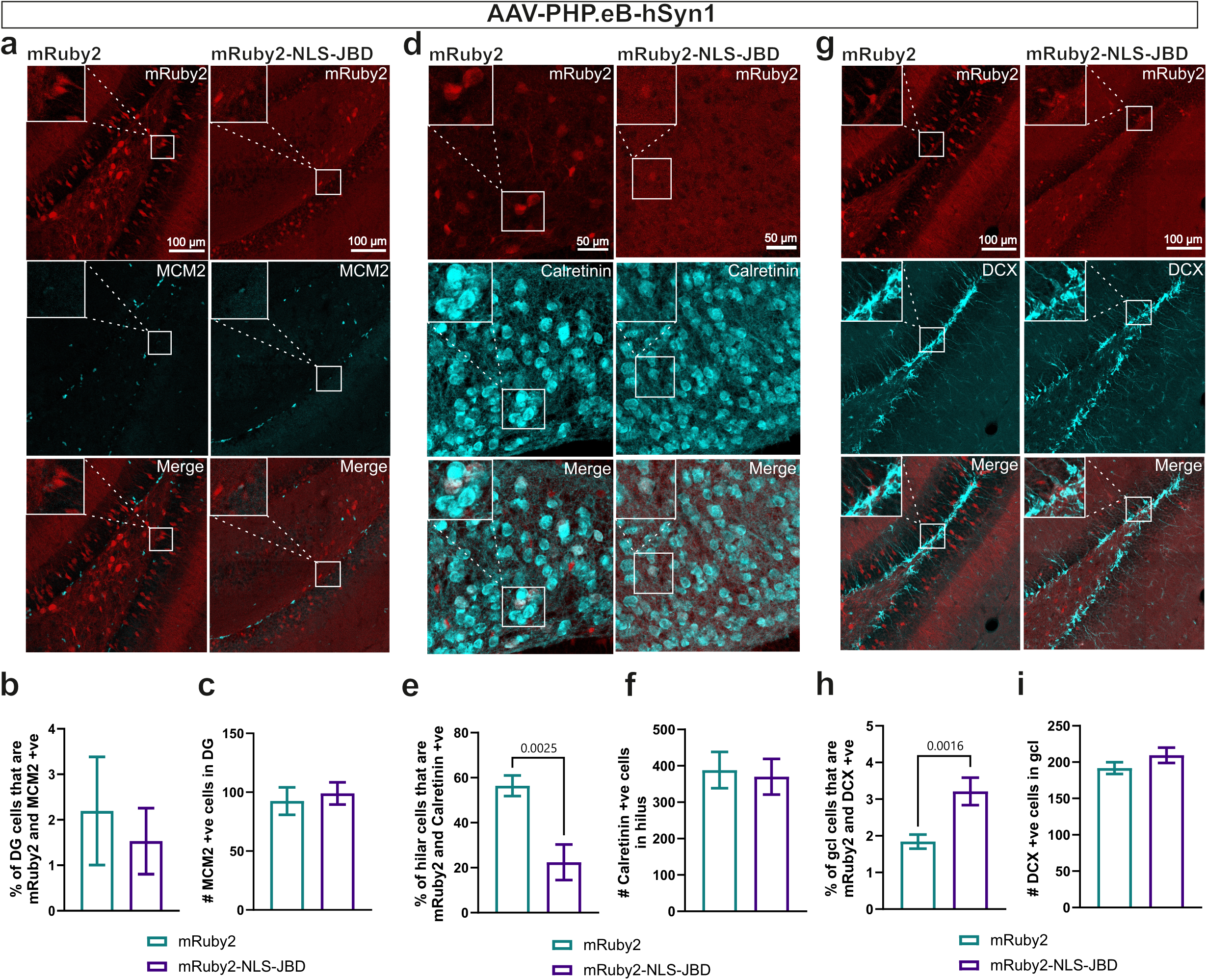
Staining of neurogenic niche cells in the ventral hippocampus dentate gyrus of AAV-PHP.eB-hSyn1-mRuby2-NLS-JBD mice with low anxiety-like phenotype. **a)** Representative maximum projection images of hSyn1-mRuby2 and hSyn1-mRuby2-NLS-JBD vector expression with MCM2 staining in the dentate gyrus (DG) of ventral hippocampus. **b)** Percentage of mRuby2 +ve cells that express MCM2 and **c)** quantification of MCM2+ve cells in the DG. Cell counting was done on a total of eight images from two hSyn1-mRuby2 mice and on a total of 12 images from four hSyn1-mRuby2-NLS-JBD mice. **d)** Representative maximum projection images of hSyn1-mRuby2 and hSyn1-mRuby2-NLS-JBD vector expression with calretinin staining in the DG hilus of ventral hippocampus. **e)** Percentage of mRuby2+ve cells that express Calretinin and **f)** quantification of calretinin +ve cells in the DG hilus. Cell counting was done on a total of nine images from two hSyn1-mRuby2 mice and on a total of 11 images from four hSyn1-mRuby2-NLS-JBD mice. **g)** Representative maximum projection images of hSyn1-mRuby2 and hSyn1-mRuby2-NLS-JBD vector expression with DCX staining in the DG of ventral hippocampus. **h)** Percentage of mRuby2 +ve cells that express DCX and **i)** quantification of DCX +ve cells in the DG granular cell layer. Cell counting was done on a total of 22 images from two hSyn1-mRuby2 mice and on a total of 18 images from four hSyn1-mRuby2-NLS-JBD mice. Statistical analyses were done with Student’s two-tailed t-test.

Specifically, the ratio of MCM2+/mRuby2+ cells was unaltered in mice expressing mRuby2-NLS-JBD compared to mRuby2 (Fig. 4b). This suggests that introducing a JNK inhibitor with the PHP.eB vector does not alter the proliferating neural stem cell pool. Moreover, we did not uncover evidence of a non-cell autonomous effect either as there was no change in the overall number of MCM2+ cells in this region (Fig. 4c). These results suggest that the anxiolytic phenotype observed in these mice (Fig. 1e), was not due to a regulation of the pool of proliferating neural progenitor cells in the ventral hippocampus.

We next examined calretinin-positive mossy cells of the hilus (Fig. 4d). These large excitatory neurons influence the number and activity of dentate gyrus granule cells by direct and indirect pathways (Scharfman & Myers, 2013; Vicidomini et al., 2020; Yeh et al., 2018). A high proportion (60%) of hilar calretinin-positive cells were mRuby2-positive (Fig. 4e), indicating that the PHP.eB-hSyn1 vector efficiently targets these cells. When we compared mice transduced with AAV-PHP.eB-hSyn1-mRuby2 and AAV-PHP.eB-hSyn1-mRuby2-NLS-JBD, the % of mRuby2+ mossy cells was dramatically reduced in the presence of NLS-JBD without an effect on the overall cell number (Fig. 4e-f).

Finally, we examined whether AAV-PHP.eB-hSyn1-NLS-JBD altered the number of immature adult born neurons of the granule cell layer by examining the proportion of mRuby2+, DCX+ cells (Fig. 4g). There was an almost two-fold increase in DCX+ adult born neurons in cells where nuclear JNK inhibitor was expressed (Fig. 4h). In line with this, there was a trend towards an increase in total DCX cell number in the dentate gyrus (Fig. 4i).

Together these results indicate that inhibition of nuclear JNK via systemic delivery of AAVs in the PHP.eB vector with Syn1 promoter increases neurogenesis in the transduced population. This correlates with increased mossy cell number. This may explain the anxiolytic behaviour evoked by AAV-PHP.eB-mRuby2-NLS-JBD-expressing mice in the elevated plus maze.

To assess if AAV-mediated expression of a JNK inhibitor would affect behavioural responses to mild stress, AAV-PHP.eB-hSyn1-mRuby2 /-NLS-JBD mice underwent a cold stress test after 12 weeks of vector expression (Supplementary Fig. 2a). As a stressor, mice were subjected to cold forced swim test as previously described (von Ziegler et al., 2022), followed 10 min later by the open field and Y-maze tests conducted consecutively. Baseline behaviour had been measured three days prior to the stress. Following stress, there was no significant behavioural change in the open field or Y-maze for any of the mice. There was however a trend towards decreased time spent in the centre following stress in mice transduced with AAV-PHP.eB-hSyn1 vectors (Supplementary figure 2b-d). However, there were no behavioural differences between control and mRuby2-NLS-JBD expressing mice after exposure to stress (Supplementary Fig. 2b-d).

### Mouse plasma EV proteins associated with metabolic process are decreased in mice displaying an anxiolytic phenotype and increased in mice displaying a depressive-like phenotype

Finally, we wanted to examine how systemic AAV-mediated expression of a JNK inhibitor would alter signalling in the periphery, and if there would be any inflammatory responses. For this we performed proteomics analysis on plasma that was collected from the mice immediately following the behavioural test battery. Extracellular vesicles (EV) were purified from the plasma as previously (Wu et al., 2025) and underwent mass spectrometry analysis in DIA mode. This analysis identified 4193 proteins including classical EV markers flotillins, CD63 and CD81, among others (Fig. 5a). The most notable overall difference was that from among the plasma EV proteins identified, 87% of those from AAV-PHP.eB-hSyn1-mRuby2-NLS-JBD transduced mice showed reduced intensity compared to mice transduced with AAV-PHP.eB-hSyn-mRuby2 (Fig. 5b).

**Figure 5.**
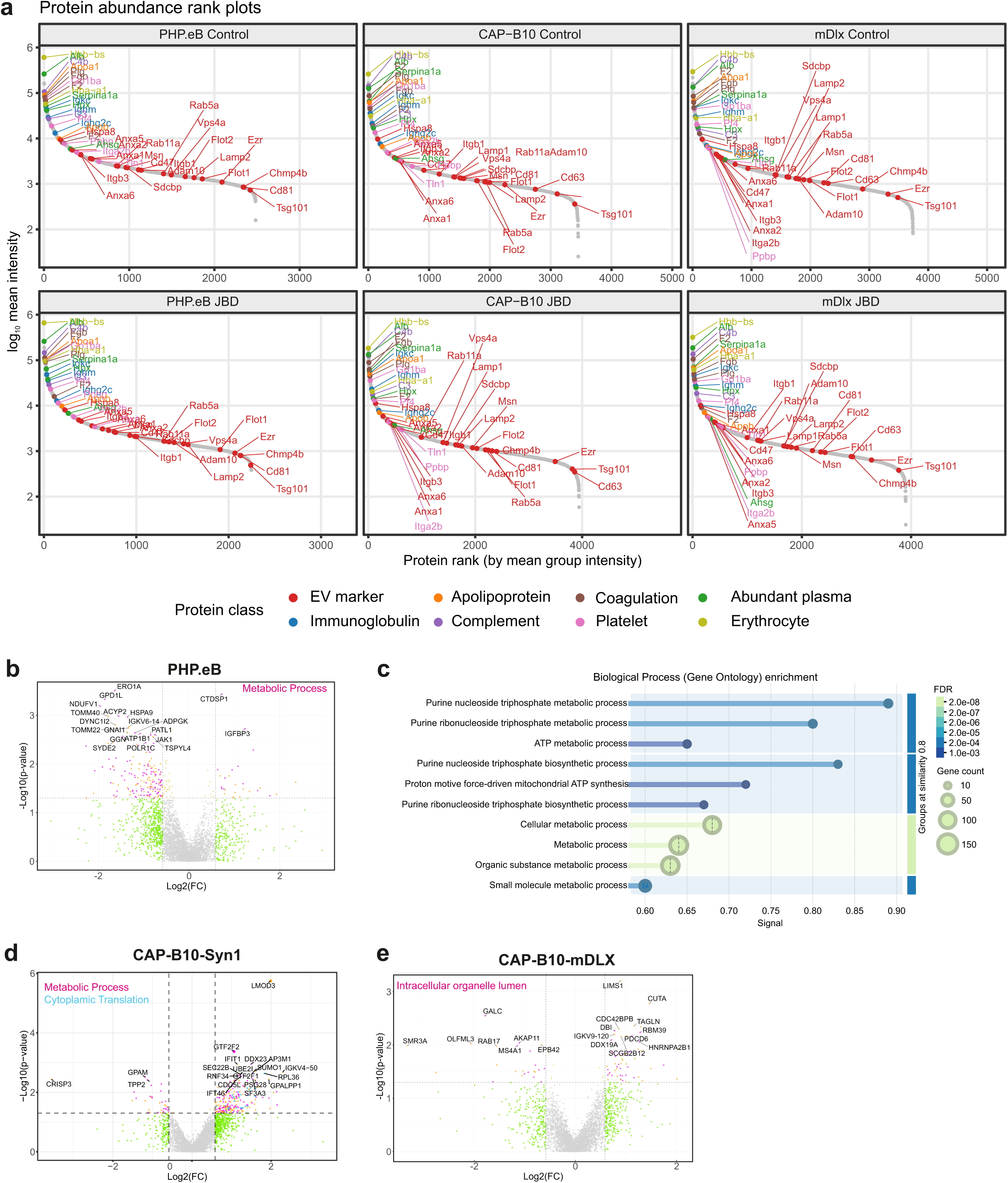
Expression of a genetically encoded JNK inhibitor in the mouse brain following systemic AAV vector administration induces protein expression changes in plasma extracellular vesicles. **a)** 12 weeks after the systemic administration of AAV vectors, plasma was collected from the mice and the extracellular vesicles purified for mass spectrometry analysis. Ranked plots show the protein classes identified in EV fractions for mice transduced with each of the vectors used in this study. EV markers are labelled in the plot if they are present in >50% of replicates. **b)** Volcano plot of significantly changing plasma EV-associated proteins from mice transduced with PHP.eB vector with or without JNK inhibitor. The top 20 most significantly changing proteins (p<0.05) are labeled. Significantly changing proteins that fall within gene ontology term metabolic process are shown in pink. **c)** The gene ontology enrichment showing the breakdown of metabolic processes that significantly change in EV proteins from AAV-PHP.eB-hSyn1-NLS-JBD mice vs control. **d)** Volcano plots from AAV.CAP-B10-hSyn1-NLS-JBD mice vs control is shown. Metabolic process proteins are marked in pink and cytoplasmic translation proteins in blue. **e)** Volcano plots from AAV.CAP-B10-mDLX-NLS-JBD mice vs control is shown. EV fraction proteins were enriched for changes in intracellular organelle lumen proteins, shown in pink. Data are from n=12 PHP-eB-hSyn1-mRuby2 mice, n=14 PHP-eB-hSyn1-mRuby2-NLS-JBD mice, n=13 CAP-B10-hSyn1-mRuby2 mice, n=13 CAP-B10-hSyn1-mRuby2-NLS-JBD mice, n=13 CAP-B10-mDlx-NLS-mRuby2 mice, n=13 CAP-B10-mDlx-NLS-mRuby2-JBD mice. Statistical analysis was done using ROTS statistical test.

Functional enrichment analysis highlighted that plasma EV cargo proteins were significantly altered in the presence of JNK inhibitor (AAV-PHP.eB-hSyn1-mRuby2-NLS-JBD). Targeted inhibition of JNK in neurons transduced by PHP.eB vector led to a decrease in proteins associated with gene ontology term metabolic process (G0008152; significance of enrichment against universal background - FDR= 9.06e-44). Metabolic process is defined as the cellular process of biochemical pathways transforming chemical substances through anabolism and catabolism (Fig. 5b, Supplementary data 1). Among the downregulated proteins were those associated with purine nucleoside triphosphate metabolism such as mitochondrial complex I proteins Ndufv1 and mt-Nd4 (Fig. 5c). In contrast, in mice transduced with the CAP-B10 plasmid with JNK inhibitor under the synapsin 1 promoter, EV associated proteins increased compared to control (Fig. 5d). These proteins were also enriched for proteins with the gene ontology term metabolic process, but also for GO term Cytoplasmic Translation. These results show that inhibition of JNK in distinct neuronal populations exerts opposing effects on the EV protein cargo in plasma. The most notable change was on proteins involved in metabolic processes.

We also examined CAP-B10-mDLX transduced mice where JNK inhibitor was targeted to GABAergic neurons. In these mice, there were no functionally enriched GO processes (Fig. 5e). Significant differences in the plasma EV pool of these mice were restricted to gene ontology terms intracellular organelle lumen (GO:0070013) and extracellular vesicles (GO: 1903561; Supplementary data 2).

These data indicate that neuronal JNK activity coordinates how metabolism-associated proteins are distributed in the circulation by plasma EVs. Moreover, it indicates that the repertoire of EV cargo is differentially regulated depending on which population of neurons in which JNK is acting, those targeted by PHP.eB-hSyn1 and CAP-B10-hSyn1 respectively.

## DISCUSSION

To determine whether the behavioural consequences of JNK inhibition depend on the cellular populations targeted, we delivered a JNK inhibitor by retro-orbital administration using three AAV capsid-promoter combinations with distinct cellular tropisms. Although all vectors exhibited widespread expression across the cortex, hippocampus, and striatum, expression from the mDlx promoter was comparatively weak. Notably, only PHP.eB-hSyn1-mediated delivery recapitulated the anxiolytic phenotype previously reported for JNK inhibition (Hong et al., 2024; Mohammad et al., 2018). In contrast, CAP-B10-mDlx-mediated expression reduced locomotor activity while enhancing novel object recognition, indicative of improved recognition memory, whereas CAP-B10-hSyn1-mediated expression increased both locomotor activity and depressive-like behaviour. Mechanistically, PHP.eB-hSyn1-mediated JNK inhibition preferentially affected hippocampal neurogenic niche populations and was associated with an increased abundance of neurogenic cells. These findings suggest that the anxiolytic effects of JNK inhibition may arise from modulation of hippocampal neurogenesis and highlight the importance of cell type-specific targeting in determining behavioural outcomes.

Surprisingly, neuron-specific expression of the JNK inhibitor under the Syn1 promoter produced an anxiolytic phenotype only when delivered using the PHP.eB capsid, whereas no comparable effect was observed following delivery with CAP-B10. One reported distinction between these vectors is that CAP-B10 exhibits a stronger preference for excitatory neurons relative to GABAergic neurons, compared with PHP.eB (Brown et al., 2021; Jang et al., 2023). However, in the present study, we did not detect obvious differences in neuronal targeting based on histological examination alone, although comprehensive cell type-specific analyses were not performed.

Notably, PHP.eB-hSyn1 transduced both hilar mossy cells and dentate granule cells. Expression of the nuclear JNK inhibitor in PHP.eB-hSyn1-treated mice was associated with a marked reduction in the number of hilar mossy cells and an approximately two-fold increase in newborn granule cells. Because hilar mossy cells form extensive connections throughout hippocampal circuitry, these changes may have important functional consequences. In the ventral dentate gyrus, mossy cells directly innervate granule cells while also promoting feed-forward inhibition of adult-born granule cells through the recruitment of local GABAergic interneurons (Galloni et al., 2022; Scharfman, 2016). Consequently, a reduction in mossy cell abundance could alter the balance of excitatory innervation and feed forward inhibition within the neurogenic niche and thereby influence neurogenesis.

Consistent with this interpretation, mice expressing mRuby2-NLS-JBD exhibited a pronounced increase in DCX-positive granule cells, indicative of enhanced neurogenesis. These findings suggest that JNK inhibition shifts cell fate dynamics within the dentate gyrus toward increased neurogenesis. Importantly, this observation is in agreement with a recent report demonstrating that selective ablation of mossy cells increases the abundance of DCX-positive neurons in the dentate gyrus (Butler et al., 2026). Together, these results raise the possibility that the anxiolytic effects observed following PHP.eB-mediated JNK inhibition are linked to remodelling of the mossy cell-neurogenesis axis within the hippocampus.

We also compared the effects of PHP.eB-hSyn1, CAP-B10-hSyn1, and CAP-B10-mDlx on the plasma extracellular vesicle (EV) proteome. Consistent with previous reports, the plasma EV proteome was enriched for proteins of hepatic origin, together with proteins derived from several other peripheral tissues (Abdelmohsen et al., 2023). The most notable difference between vector groups was observed in mice transduced with PHP.eB-hSyn1. In these animals, several brain-derived proteins, including Syntaxin-1B (Stx1b), Kif1a, Kcc2 (Slc12a5), and the brain-enriched glucose transporter Glut1, were detected in plasma EVs, whereas these proteins were not detected in EVs from CAP-B10-transduced mice. One possible explanation for this difference is the greater propensity of AAV-PHP.eB to induce astrocytic activation, which has previously been reported following systemic PHP.eB administration (Chan et al., 2017a; Rincon et al., 2018). Activated astrocytes may enhance EV release and facilitate the transfer of brain-derived vesicular cargo to the circulation (Dickens et al., 2017). Consistent with this hypothesis, all of these brain-derived proteins (Stx1b, Kif1a and Kcc2) were significantly reduced in EVs isolated from mice expressing the JNK inhibitor. Given the well-established anti-inflammatory effects of JNK inhibition (Mielke & Herdegen, 2000), these findings suggest that inflammatory processes may contribute to the appearance of brain-derived proteins in circulating EVs.

Among the JNK-inhibitor regulated proteins of particular interest was Stx1b, which plays an essential role in synaptic vesicle exocytosis and neurotransmitter release (Jahn et al., 2023). In a recent phosphoproteomic study from our laboratory, Stx1b emerged as a central hub within a signalling network associated with anxiety-like behaviour (Hong et al., 2026).

Notably, decreased phosphorylation of Stx1b at Ser14 was associated with reduced anxiety-like behaviour in that study. In the present work, Stx1b abundance in plasma EVs was reduced by approximately two-fold in mice expressing the JNK inhibitor. Although the relationship between EV-associated Stx1b abundance and neuronal Stx1b regulation remains unclear, the convergence of these findings implicates Stx1b-regulated synaptic signalling in the anxiety-like behaviour controlled by JNK.

Taken together, the increased representation of brain-derived proteins in plasma EVs from PHP.eB-transduced mice is consistent with enhanced neuroinflammatory or glial responses, whereas the reduction of these proteins following JNK inhibition is compatible with the anti-inflammatory actions of JNK pathway suppression in neurons. These findings suggest that circulating EVs may provide a peripheral readout of brain-specific molecular processes induced by AAV-mediated gene delivery and JNK signalling.

Changes in several proteins associated with inflammatory signaling were observed in PHP.eB mice. These included angiopoietin-1, β-catenin, Janus kinase 1 (JAK1), lactate dehydrogenase B (LDHB), and 14-3-3γ, consistent with previous reports linking these pathways to JNK activity (Hong et al., 2024; Tsuruta et al., 2004). The overall protein profile (including the unchanged/increased angiopoietin-1) is consistent with an anti-inflammatory effect.

LDHB is generally considered a non-specific marker of tissue damage and cell death. Its reduction may reflect diminished apoptotic signalling following JNK inhibition, as JNK promotes AP-1-dependent transcriptional programs associated with apoptosis and the *Ldhb* promoter contains several AP-1 binding sites (Semba et al., 2020). Alternatively, decreased LDHB expression may reflect altered lactate metabolism and a shift toward aerobic glycolysis (the Warburg effect), a metabolic phenotype previously associated with JNK inhibition (Hong et al., 2026; Luo et al., 2026).

Proteins involved in metabolic processes were among the most strongly affected components of the plasma EV proteome following neuronal expression of the JNK inhibitor. These proteins were predominantly reduced in mice transduced with AAV-PHP.eB-hSyn1-JBD, whereas they were increased in mice receiving AAV-CAP-B10-hSyn1-JBD. The opposite directionality of these changes may reflect differences in the neuronal populations transduced by the two vectors, suggesting that distinct neurocircuits contribute differently to the regulation of peripheral metabolism. Nevertheless, the prominence of metabolic pathways in both datasets is consistent with the established role of JNK as a central regulator of metabolic homeostasis. JNK signaling contributes to metabolic adaptations induced by inflammation and oxidative stress, and its inhibition can modify multiple metabolic pathways (Fernández-González et al., 2026).

The widespread changes in metabolic proteins are unlikely to result from direct inhibition of JNK in peripheral tissues, as expression of the transgene is driven by the neuron-specific Synapsin1 promoter. Instead, they may reflect central regulation of peripheral metabolism, as JNK is known to influence metabolic processes through hypothalamic neuroendocrine and autonomic pathways (Belgardt et al., 2010; Fernández-González et al., 2025; Sabio et al., 2010). These findings therefore support the existence of molecular communication between the brain and the periphery. However, a contribution from the enteric nervous system cannot be excluded, as Synapsin1 is also expressed in neurons of the gastrointestinal tract.

Furthermore, both PHP.eB and CAP-B10 capsids are detected only at low levels in peripheral tissues such as the liver relative to the CNS (Finneran et al., 2021; Goertsen et al., 2021; Jackson et al., 2016).

The increased transfer of CNS-derived EVs to the periphery has been reported in response to diverse physiological and pathological stimuli, including nutritional perturbations, metabolic changes, and cellular stress, all processes closely associated with JNK signalling (Fernández-González et al., 2025). Consistent with this, neuronal JNK inhibition altered peripheral inflammatory and metabolic signatures despite largely CNS-restricted transgene expression, supporting a role for brain-to-periphery communication in mediating known systemic effects of JNK signalling. Most importantly, however, only AAV-PHP.eB-hSyn1-mediated delivery of the JNK inhibitor reproduced the previously reported anxiolytic phenotype. The absence of a comparable effect following CAP-B10-hSyn1 or CAP-B10-mDlx delivery indicates that the behavioural consequences of JNK inhibition depend critically on the neuronal populations targeted. Our results suggest that anxiety-related behaviour is regulated by specific JNK-sensitive neural circuits in the hippocampal neurogenic niche rather than by global neuronal JNK activity and demonstrate the value of cell type-selective AAV targeting for dissecting the neural substrates of complex behaviours.

## MATERIALS & METHODS

### Plasmid construction

NLS-JBD was cut from pEGFP-NLS-JBD (Björkblom et al., 2005) and ligated into the BsrGI and HindIII sites of pAAV-CAG-mRuby2 (Addgene plasmid # 99123 ; http://n2t.net/addgene:99123 ; RRID:Addgene_99123) and pAAV-hSyn1-mRuby2 (Addgene plasmid # 99126 ; http://n2t.net/addgene:99126 ; RRID:Addgene_99126) to yield pCAG-mRuby2-NLS-JBD and phSyn1-mRuby2-NLS-JBD, respectively. pCAG-mRuby2 and phSyn1-mRuby2 were used as the control plasmids. To generate pAAV-mDlx-NLS-mRuby2-JBD, the JBD sequence was ligated into the BsrGI and AscI sites of the control plasmid pAAV-mDlx-NLS-mRuby2 (Addgene plasmid # 99130 ; http://n2t.net/addgene:99130 ; RRID:Addgene_99130).

### AAV production

Viral vectors were produced as previously described (Challis et al., 2019). Briefly, 80-90 % confluent HEK293T cells grown in Dulbecco’s modified Eagle’s medium (DMEM) supplemented with 5 % FBS were triple-transfected with a helper plasmid (Addgene 112867), a packaging plasmid (PHP.eB: Addgene 103005; CAP-B10: Addgene, 175004), and a plasmid containing the transgene in a ratio of 2:4:1, respectively, using polyethylenimine. Media was replaced with fresh DMEM 24 hours post-transfection, and collected at 72 hours and 120 hours post-transfection. Cells were harvested at 120 hours and pelleted with centrifugation (2000 g, 15 min, RT), lysed with repeated freeze-thaw cycles in lysis buffer (0.001% pluronic F68 + 200mM NaCl + 1 mM MgCl2 in PBS) and cleared from cell debris by centrifugation (3200 g, 15 min, 4 °C). The media collected from cells was centrifuged (2000 g, 15 min, RT) and filtered through a 0.45 µm PES membrane, followed by overnight AAV precipitation with PEG at 4 °C. The precipitated AAVs were pelleted by centrifugation (2820 g, 15 min, 4 °C) and resuspended in lysis buffer. The resuspended viruses were combined with the supernatant containing viruses from cells and incubated with 50 units of benzonase per ml of viral suspension for 45 min at 37 °C, followed by centrifugation (2415 g, 10 min, 4 °C). Clarified supernatant was loaded on top of iodixanol step gradient (15%-60%) and ultracentrifuged (350 000 g, 90 min, 10 °C), after which the viruses were collected from the 40 % iodixanol layer. To remove iodixanol and concentrate the viruses, the Amicon filter device (Amicon® Ultra Centrifugal Filter, 100 kDa MWCO, #UFC9100 Millipore) membrane was first primed by incubation with 15 ml of 0.1 % Pluronic-F68 in PBS for 10 min at RT, followed by centrifugation (3000 g, 5 min, °C) with 15 ml of 0.01 % Pluronic-F68 in PBS and then centrifugation (3000 g, 5 min, 4 °C) with 15 ml of 0.001 % Pluronic-F68 in PBS with 200 mM NaCl. The virus sample was then added onto the membrane with 0.001 % Pluronic-F68 and centrifuged (3000 g, 4 min, 4°C). This was repeated until all of the viral solution was centrifuged with the filter membrane. The final vector solution was aliquoted, snap-frozen with liquid nitrogen and stored at -80 °C until use. Viral titer was determined by quantitative PCR.

### Animals and intravenous delivery of AAV vectors

Adult 8-wk-old C57BL/6J male mice (Charles River, Germany) were maintained at the Central Animal Laboratory of the University of Turku with ambient temperature of 21 ± 3 °C and humidity of 55 ± 15 % on a 12 h light/dark cycle with food and water ad libitum. Total number of mice injected with AAV vectors and used in behavioural testing was 12-14 per vector. Detailed animal numbers of each group are indicated in the figure legends.

Procedures and behavioural tests were performed between 8.00 and 15.00. Behavioural tests were conducted every other day with one day of rest in between. Animals were allowed to acclimatize to the room for 1 h before behavioural testing. Arenas and objects therein used for testing were cleaned using 70 % EtOH between each animal. The experimenter was blind to the viral expression throughout the experiments. Procedures and behavioural tests were approved by the National Animal Experiment Board ELLA (licenses ESAVI/16343/2019 and ESAVI/30686/2022).

The AAV vectors were intravenously injected into the retro-orbital sinus of the mice following anaesthesia induction with 2.5 % isoflurane, maintained with 2 % isoflurane on a heating pad. Oxybuprocaine (Oftan Obucain 4 mg/ml, Santen Oy, Tampere, Finland), an ophthalmic anaesthetic, was applied to the right eye before injection. Viscotears gel (Novartis, Basel, Switzerland) was applied to the left eye to prevent dryness during anaesthesia. The needle was introduced into the retro-orbital sinus through the medial canthus and 1×10^7^ TU of the AAV-PHP.eB-CAG vectors, 1×10^10^ vg of the AAV-PHP.eB-hSyn1 vectors, 3×10^11^ vg of the AAV.CAP-B10-hSyn1 vectors, or 3×10^11^ vg of the AAV.CAP-B10-mDlx vectors diluted in sterile PBS in a final volume of 40 µl was injected slowly at the base of the eye. After injection, mice were returned to their home cages on a heating pad until recovery from anaesthesia. Mice were kept single-housed after viral injection.

### Behaviour tests

6 wk after AAV injection, mice were tested in the open field, novel object recognition, elevated plus maze, Y-maze, and forced swim tests in the order described below. For tracking the mice and behaviour analysis, the EthoVision XT 13 video tracking system (Noldus Information Technology, Wageningen, The Netherlands) with the Basler ace acA1300-60gc (Basler AG, Ahrensburg, Germany) camera was used. Tracking of animals and behaviours was done from the center point of the mice.

### Open field

Mice were placed in the corner of the 45 x 45 x 45 cm square arena (made from white plastic) facing the center and their activity was recorded for 10 min. During Ethovision analysis, the arena was divided into the center zone (22.5 x 22.5 cm) and the peripheral zone. Total distance moved, mean velocity, number of entries in the center, cumulative duration in the center, and latency to the first entry to the center were scored by the EthoVision software.

### Novel object recognition

The novel object recognition test was conducted in the same arena as open field and the test consisted of three phases (habituation, training, and testing phase) all of which were done during the same day. In the habituation phase, the mouse was placed in an empty arena for 2 min, after which the mouse was then put back in its home cage. After 1 min, in the training phase, the mouse was placed in the arena with two identical objects in adjacent corners 8 cm off from walls and left to explore for 5 min or until 30 sec of object exploration was recorded. The mouse was then put back in its home cage for two hours. In the testing phase, the mouse was placed in the arena with one familiar object from the training phase and one novel object and left to explore for 5 minutes. In all phases, mice were placed in the arena in the lower corner facing the center of the arena.

Objects used in the test were 5 x 2,7 x 9 cm culture flasks filled with grey sand as familiar objects and a 6,5 x 3 x 8 cm block made of four different coloured Lego bricks as novel objects. The exploration time of each object was manually scored from videos by the experimenter. Behaviours were considered explorative if the full attention of the mouse was towards the object i.e., sniffing and touching the object, at a close distance. Analyses were made from the testing phase. Time spent exploring the familiar and novel objects, total time spent exploring both objects, discrimination index, and novel preference index were measured. The discrimination index was calculated as (TIME NOVEL – TIME FAMILIAR) / TOTAL TIME EXPLORING BOTH OBJECTS. Novel preference index was calculated as TIME NOVEL/TOTAL TIME EXPLORING BOTH OBJECTS.

### Elevated plus maze

Mice were placed in the center of a plus maze with two open (35 x 5 cm) and two closed (35 x 5 x 15 cm) arms elevated at 50 cm above the floor, facing an open arm. Behaviour was recorded for 5 min. Total distance moved, mean velocity, cumulative duration spent in the open and closed arms of the maze, number of entries into the arms, and latency to enter the arms were scored by the EthoVision software.

### Y-maze

Mice were placed at the end of one arm of the Y-shaped arena with three arms (30 x 5 x 15 cm) facing the center and left to explore for 10 min. Alternations and max alternations were scored by the EthoVision software. Spontaneous alternation percentage was calculated as (ALTERNATIONS / MAX ALTERNATIONS) * 100.

### Forced swim test

Mice were placed in an 18 cm diameter glass beaker with 24 ± 1°C water at 15 cm from the bottom of the beaker for 6 minutes. The activity was recorded using a Samsung SMX-F300BP camera (Suwon, South Korea). Time spent immobile and latency to immobility were manually scored by the experimenter. Latency to immobility was scored from the six-minute recording whereas time spent immobile was scored from the last four minutes of the recording. The mice were considered to be immobile if they were motionless or had only small movements to keep their balance. After the test, the mice were dried with towels and returned to their home cages over heating pads.

### Stress paradigm

12 weeks (PHP.eB-hSyn1 mice) or 9 weeks (CAP-B10-hSyn1 and mDlx mice) after AAV injection, the mice that underwent behavioural testing were subjected to a stress test protocol to assess the effects of stress and JNK inhibition on behaviour. Baseline behaviour of the mice was tested with the open field and Y-maze tests consecutively in the same day. Three days after baseline measurement, the mice were first subjected to cold forced swim test as described previously (von Ziegler et al., 2022) to induce stress. In brief, the test was conducted the same as described above but with 18 ± 0.1°C water temperature. After the forced swim test, the mice were returned to their home cage on a heating pad for ten minutes, after which the open field and Y-maze tests were conducted consecutively. PHP.eB underwent testing at baseline and after stress exposure, whereas CAP-B10-hSyn1 and mDlx mice were additionally measured one day after stress exposure.

### Perfusion and tissue processing

For collection of brain tissue for immunofluorescence analysis, mice were terminally by intraperitoneal injection with sodium pentobarbital (30 mg ml^−1^ Mebunate, Orion Pharma, Espoo, Finland) followed by intracardial perfusion with 4 % paraformaldehyde (PFA) in PBS. Brains of the mice were collected and post-fixed in 4 % PFA in PBS for 24 hours at 4 °C. Next, the brains were moved to 30 % sucrose in PBS overnight or until the brain had sunk to the bottom of the solution. The brains were then snap-frozen in isopentane (Sigma-Aldrich, St. Louis, USA) chilled with dry ice and stored at -80 °C until use. Before sectioning, the brains were kept at -20 °C overnight. The brains were sectioned using a cryostat (CM1950, Leica Biosystems, Nussloch, Germany) into 40 µm coronal and sagittal sections and kept in PBS for staining and storing.

For collection of plasma after the stress test paradigm, mice were immediately sacrificed by cervical dislocation and the blood was drained by making an incision on the neck. The blood was collected into Eppendorf tubes with a drop of 0.5 M EDTA. The tubes were kept on ice until centrifugation at 5000 RPM for 10 min. The plasma was then collected, snap-frozen and stored at -80 °C.

### Immunofluorescence staining and imaging

Free-floating brain sections were washed 3 x 5 min with PBS and incubated in a blocking solution (PBS containing 1 % bovine serum albumin and 0.4 % Triton X-100) for 1 hour at room temperature. Sections were then incubated with primary antibodies in the blocking solution for 72 hours at 4 °C on gentle shaking. Next, sections were washed 3 × 10 min with PBS before adding secondary antibodies with 1:1000 Hoechst-33342 in blocking solution and incubated for 2 hours.

Sections were then washed 3 × 10 min with PBS and transferred onto slides and left to dry overnight. Coverslips were mounted using Mowiol mounting medium (Sigma-Aldrich).

Primary antibodies used were 1:1000 goat anti-DCX (sc-8066, Santa Cruz Biotechnology, Dallas, TX, USA), 1:500 rabbit anti-MCM2 (ab4461, Abcam, Cambridge, United Kingdom), and 1:1000 mouse anti-Calretinin (610908, BD Transduction Laboratories, Franklin Lakes, NJ, USA). For detection, secondary antibodies conjugated to Alexa-488 (A-11055, A-21206, A-21202, Invitrogen, Waltham, Massachusetts, U.S.) were used. mRuby2 signal from AAV-transduced cells was visualized from the native signal without staining.

Tiled whole-brain images of mRuby2 signal were acquired with the THUNDER Imager Live Cell (Leica Microsystems, Wetzlar, Germany) widefield microscope with a 10x objective (0.45 NA) using 554 nm excitation and 594 nm emission. Imaging of the immunofluorescence cell type markers was done using the Zeiss LSM880 (Zeiss, Oberkochen, Germany) confocal microscope with a 10x objective (0.3 NA) using 488 nm excitation and 530 nm emission for Alexa-488 and 543 nm excitation and 633 nm emission with for mRuby2. In the analysis, the number of cells expressing the markers were calculated from maximum projection images of 10 z-sections of 1.79 µm optical section each.

For imaging of mRuby2 expression in the brain, 4 brains from each group were imaged, with the exception of 2 brains from the AAV-PHP.eB-hSyn1-mRuby2 group and 2 brains from the AAV.CAP-B10-mDlx-NLS-mRuby2 /-JBD groups. For immunofluorescence cell type analysis, 2 brains from the AAV-PHP.eB-hSyn1-mRuby2 and 4 brains from AAV-PHP.eB-hSyn1-mRuby2-NLS-JBD groups were used. DCX, MCM2, and calretinin cell counts were performed from 22, 8, and 9 AAV-PHP.eB-hSyn1-mRuby2 images and 18, 12, and 11 AAV-PHP.eB-hSyn1-mRuby2-NLS-JBD images, respectively.

### Plasma EV purification and MS

*Sample digestion* – Frozen plasma samples were thawed at room temperature. After thawing, Bis Tris propane solution and sodium chloride were added to 20 µl of each sample to reach final concentrations of 50 mM Bis Tris propane (pH 6.3) and 150 mM sodium chloride. The samples were then brought to a final volume of 50 µl using wash buffer, consisting of 50 mM Bis Tris propane with 150 mM sodium chloride at pH 6.3. Sample preparation was performed using an Opentrons OT 2 liquid-handling robot (Opentrons Labworks, Long Island City, NY, USA) using a Magnet procol (Wu et al., 2025) following modifications and automatization made available by Evosep Biosystems (Evosep Biosystems, Odense, Denmark). These Evosep provided modifications were implemented on the OT 2 to enable fully automated bead-based sample processing.

Plasma samples were mixed five times with 1 µl of MagReSyn strong anion-exchange beads (ReSyn Biosciences, Edanvale, Gauteng, South Africa) in a 96 well plate to facilitate protein binding. The beads were then washed three times with wash buffer. Protein extraction and alkylation were carried out in 50 mM ammonium bicarbonate containing 1% sodium dodecyl sulfate, 5 mM tris(2 carboxyethyl)phosphine hydrochloride, and 10 mM chloroacetamide for 1 h with mixing.

Following extraction, acetonitrile (ACN) was added to a final concentration of 70%, and samples were incubated with mixing for two consecutive 10 min periods. The ACN solution was then replaced once with 100% ACN to remove impurities and removed for protein digestion. Proteins were digested for 16 h at 37 °C in a humidified chamber using 50 mM ammonium bicarbonate supplemented with 300 ng trypsin and 75 ng Lys C per sample. After digestion, samples were acidified with 0.1% formic acid, and 50% of each sample was loaded onto Evotip Pure tips (Evosep Biosystems) using the Opentrons OT 2 platform.

### MS analysis

Samples were analysed by data-independent acquisition (DIA) mass spectrometry in diaPASEF mode (Meier et al., 2020) with timsTOF fleX MALDI-2 (Bruker Daltonics, Billerica, MA, USA) in electrospray ionisation mode. The mass spectrometer was interfaced with an Evosep One (Evosep Biosystems) operated with the 30 samples per day method, using an Evosep Endurance column (EV1106; 15 cm × 150 µm, 1.9 µm ReproSil-Pur C18). Peptides were ionised using a CaptiveSpray 2 source (Bruker Daltonics) at a capillary voltage of 1500 V, with dry gas at 3.0 L/min and 180 °C. Data were acquired in positive ion mode over an *m/z* range of 100–1700 and an ion mobility (1/K0) range of 0.60– 1.60 V·s·cm⁻². Ion accumulation and ramp times were both 100 ms (100% duty cycle). Each acquisition cycle consisted of one MS1 frame followed by 25 diaPASEF frames, each containing two variable-width isolation windows (50 windows in total, 7–50 Th), together covering *m/z* 400–1099 and 1/K0 0.70–1.30 V·s·cm⁻², resulting in a cycle time of 2.77 s. Ion mobility was calibrated using three ions of the Agilent ESI-L Tuning Mix (*m/z* 622, 922, 1222).

### MS data quantification

Raw data were processed in Spectronaut 19 (Biognosys, Switzerland) using directDIA against the UniProtKB Mus musculus proteome (Swiss-Prot and TrEMBL, including isoforms; accessed 18 November 2024), with default settings unless otherwise stated and imputation disabled. Carbamidomethylation of cysteine was set as a fixed modification, and protein N-terminal acetylation and methionine oxidation as variable modifications. Global median normalisation was applied, protein quantities were calculated using MaxLFQ, and deepQuant was enabled. Protein intensities were log2-transformed, and proteins quantified in at least 50% of samples in at least one comparison group were retained. Remaining missing values were imputed in Perseus 1.6.15.0 (Tyanova et al., 2016) from a normal distribution based on the total matrix with default settings. Differential abundance was assessed with the Reproducibility-Optimized Test Statistic as implemented in PhosPiR (Hong et al., 2022). Plots were generated with ggplot2 in R.

### Statistical analysis

To test for statistical significance between two groups, an unpaired Student’s t-test was used. For analysis of OF distance moved in time bins, two-way ANOVA was used. Statistical analysis and graphs were done using the GraphPad Prism version 8 for Windows (GraphPad Software, San Diego, California USA, www.graphpad.com). The graphs show the mean with ± SEM of the groups. A p-value <0.05 was considered significant, and nonsignificant differences were not marked on the graphs.

## ACKNOWLEDGEMENTS

This work was supported by grants from the Research Council of Finland #334776 #362838 #379008, Turku Doctoral Programme of Molecular Medicine, the Finnish Brain Foundation, and the Otto A. Malm’s Donation Fund.

## REFERENCES

Abdelmohsen, K., Herman, A. B., Carr, A. E., Henry-Smith, C. A., Rossi, M., Meng, Q., Yang, J. H., Tsitsipatis, D., Bangura, A., Munk, R., Martindale, J. L., Nogueras-Ortiz, C. J., Hao, J., Gong, Y., Liu, Y., Cui, C. Y., Hartnell, L. M., Price, N. L., Ferrucci, L., … Gorospe, M. (2023). Survey of organ-derived small extracellular vesicles and particles (sEVPs) to identify selective protein markers in mouse serum. Journal of Extracellular Biology, 2(8), e106. 10.1002/JEX2.106

Belgardt, B. F., Mauer, J., Wunderlich, F. T., Ernst, M. B., Pal, M., Spohn, G., Brönneke, H. S., Brodesser, S., Hampel, B., Schauss, A. C., & Brüning, J. C. (2010). Hypothalamic and pituitary c-Jun N-terminal kinase 1 signaling coordinately regulates glucose metabolism. Proceedings of the National Academy of Sciences of the United States of America, 107(13), 6028. 10.1073/PNAS.1001796107

Björkblom, B., Östman, N., Hongisto, V., Komarovski, V., Filén, J. J., Nyman, T. A., Kallunki, T., Courtney, M. J., & Coffey, E. T. (2005). Constitutively Active Cytoplasmic c-Jun N-Terminal Kinase 1 Is a Dominant Regulator of Dendritic Architecture: Role of Microtubule-Associated Protein 2 as an Effector. The Journal of Neuroscience, 25(27), 6350. 10.1523/JNEUROSCI.1517-05.2005

Brown, D., Altermatt, M., Dobreva, T., Chen, S., Wang, A., Thomson, M., & Gradinaru, V. (2021). Deep Parallel Characterization of AAV Tropism and AAV-Mediated Transcriptional Changes via Single-Cell RNA Sequencing. Frontiers in Immunology, 12, 730825. 10.3389/FIMMU.2021.730825/FULL

Butler, C. R., Squellati, C. R., Danis, A. B., Isakharov, A., Westbrook, G. L., & Schnell, E. (2026). Hilar mossy cells control structural and functional organization in the dentate gyrus. bioRxiv, 2026.07.15.738260. 10.64898/2026.07.15.738260

Byrne, B. J., Flanigan, K. M., Matesanz, S. E., Finkel, R. S., Waldrop, M. A., D’Ambrosio, E. S., Johnson, N. E., Smith, B. K., Bönnemann, C., Carrig, S., Rossano, J. W., Greenberg, B., Lalaguna, L., Lara-Pezzi, E., Subramony, S., Corti, M., Mercado-Rodriguez, C., Leon-Astudillo, C., Ahrens-Nicklas, R., … George, L. A. (2025). Current clinical applications of AAV-mediated gene therapy. Molecular Therapy, 33(6), 2479. 10.1016/J.YMTHE.2025.04.045

Challis, R. C., Ravindra Kumar, S., Chan, K. Y., Challis, C., Beadle, K., Jang, M. J., Kim, H. M., Rajendran, P. S., Tompkins, J. D., Shivkumar, K., Deverman, B. E., & Gradinaru, V. (2019). Systemic AAV vectors for widespread and targeted gene delivery in rodents. Nature Protocols 2019 14:2, 14(2), 379–414. 10.1038/s41596-018-0097-3

Chan, K. Y., Jang, M. J., Yoo, B. B., Greenbaum, A., Ravi, N., Wu, W. L., Sánchez-Guardado, L., Lois, C., Mazmanian, S. K., Deverman, B. E., & Gradinaru, V. (2017a). Engineered AAVs for efficient noninvasive gene delivery to the central and peripheral nervous systems. Nature Neuroscience, 20(8), 1172. 10.1038/NN.4593

Chan, K. Y., Jang, M. J., Yoo, B. B., Greenbaum, A., Ravi, N., Wu, W.-L., Sánchez-Guardado, L., Lois, C., Mazmanian, S. K., Deverman, B. E., & Gradinaru, V. (2017b). Engineered AAVs for efficient noninvasive gene delivery to the central and peripheral nervous systems. Nature Neuroscience, 20(8), 1172. 10.1038/NN.4593

Corrales, T. de los R., Losada-Pérez, M., & Casas-Tintó, S. (2021). JNK Pathway in CNS Pathologies. International Journal of Molecular Sciences, 22(8). 10.3390/IJMS22083883

David, D. J., Samuels, B. A., Rainer, Q., Wang, J. W., Marsteller, D., Mendez, I., Drew, M., Craig, D. A., Guiard, B. P., Guilloux, J. P., Artymyshyn, R. P., Gardier, A. M., Gerald, C., Antonijevic, I. A., Leonardo, E. D., & Hen, R. (2009). Neurogenesis-Dependent and -Independent Effects of Fluoxetine in an Animal Model of Anxiety/Depression. Neuron, 62(4), 479–493. 10.1016/j.neuron.2009.04.017

Dickens, A. M., Tovar-Y-Romo, L. B., Yoo, S. W., Trout, A. L., Bae, M., Kanmogne, M., Megra, B., Williams, D. W., Witwer, K. W., Gacias, M., Tabatadze, N., Cole, R. N., Casaccia, P., Berman, J. W., Anthony, D. C., & Haughey, N. J. (2017). Astrocyte-shed extracellular vesicles regulate the peripheral leukocyte response to inflammatory brain lesions. Science Signaling, 10(473), eaai7696. 10.1126/SCISIGNAL.AAI7696

Fernández-González, I., Freire-Agulleiro, O., Ferreira, V., Silveira-Loureiro, M., Rial-Pensado, E., Garrido-Gil, P., Martínez, G., Rada, P., Labandeira-García, J. L., Mittag, J., González-García, I., Diéguez, C., Nogueiras, R., Valverde, Á. M., Davis, R. J., Sabio, G., Barca-Mayo, O., & López, M. (2025). JNK1 in SF1 neurons regulates the central action of thyroid hormones on hepatic lipid metabolism. Molecular Metabolism, 98, 102170. 10.1016/J.MOLMET.2025.102170

Fernández-González, I., Vattathara, J. J., Davis, R. J., Sabio, G., & López, M. (2026). Attack of the kinases: JNK signaling in metabolism. American Journal of Physiology. Regulatory, Integrative and Comparative Physiology, 330(4), R379–R395. 10.1152/AJPREGU.00286.2025

Finneran, D. J., Njoku, I. P., Flores-Pazarin, D., Ranabothu, M. R., Nash, K. R., Morgan, D., & Gordon, M. N. (2021). Toward Development of Neuron Specific Transduction After Systemic Delivery of Viral Vectors. Frontiers in Neurology, 12, 685802. 10.3389/FNEUR.2021.685802/TEXT

Galeotti, N., & Ghelardini, C. (2012). Regionally selective activation and differential regulation of ERK, JNK and p38 MAP kinase signalling pathway by protein kinase C in mood modulation. The International Journal of Neuropsychopharmacology, 15(6), 781–793. 10.1017/S1461145711000897

Galloni, A. R., Samadzelkava, A., Hiremath, K., Oumnov, R., & Milstein, A. D. (2022). Recurrent Excitatory Feedback From Mossy Cells Enhances Sparsity and Pattern Separation in the Dentate Gyrus via Indirect Feedback Inhibition. Frontiers in Computational Neuroscience, 16, 826278. 10.3389/FNCOM.2022.826278/FULL

Goertsen, D., Flytzanis, N. C., Goeden, N., Chuapoco, M. R., Cummins, A., Chen, Y., Fan, Y., Zhang, Q., Sharma, J., Duan, Y., Wang, L., Feng, G., Chen, Y., Ip, N. Y., Pickel, J., & Gradinaru, V. (2021). AAV capsid variants with brain-wide transgene expression and decreased liver targeting after intravenous delivery in mouse and marmoset. Nature Neuroscience 2021 25:1, 25(1), 106–115. 10.1038/s41593-021-00969-4

Hong, Y., Flinkman, D., Suomi, T., Pietilä, S., James, P., Coffey, E., & Elo, L. L. (2022). PhosPiR: an automated phosphoproteomic pipeline in R. Briefings in Bioinformatics, 23(1). 10.1093/BIB/BBAB510

Hong, Y., Siino, V., Flinkman, D., Deshpande, P., Martinez, S. O., Fagerholm, V., Varidaki, A., Heemeryck, P., Sourander, C., James, P., & Coffey, E. (2026). JNK-regulated phosphoproteome links synaptic and metabolic pathways to mood regulation. Neurobiology of Disease, 218, 107207. 10.1016/J.NBD.2025.107207

Hong, Y., Sourander, C., Hackl, B., Patton, J. S., John, J., Paatero, I., & Coffey, E. (2024). Jnk1 and downstream signalling hubs regulate anxiety-like behaviours in a zebrafish larvae phenotypic screen. Scientific Reports, 14(1). 10.1038/S41598-024-61337-3

Jackson, K. L., Dayton, R. D., Deverman, B. E., & Klein, R. L. (2016). Better Targeting, Better Efficiency for Wide-Scale Neuronal Transduction with the Synapsin Promoter and AAV-PHP.B. Frontiers in Molecular Neuroscience, 9(NOV2016). 10.3389/FNMOL.2016.00116

Jahn, R., Cafiso, D. C., & Tamm, L. K. (2023). Mechanisms of SNARE proteins in membrane fusion. Nature Reviews. Molecular Cell Biology, 25(2), 101. 10.1038/S41580-023-00668-X

Jang, M. J., Coughlin, G. M., Jackson, C. R., Chen, X., Chuapoco, M. R., Vendemiatti, J. L., Wang, A. Z., & Gradinaru, V. (2023). Spatial transcriptomics for profiling the tropism of viral vectors in tissues. Nature Biotechnology, 41(9), 1272. 10.1038/S41587-022-01648-W

Kuan, C. Y., Yang, D. D., Samanta Roy, D. R., Davis, R. J., Rakic, P., & Flavell, R. A. (1999). The Jnk1 and Jnk2 protein kinases are required for regional specific apoptosis during early brain development. Neuron, 22(4), 667–676. 10.1016/S0896-6273(00)80727-8

Kügler, S., Kilic, E., & Bähr, M. (2003). Human synapsin 1 gene promoter confers highly neuron-specific long-term transgene expression from an adenoviral vector in the adult rat brain depending on the transduced area. Gene Therapy, 10(4), 337–347. 10.1038/SJ.GT.3301905

Li, X. M., Li, C. C., Yu, S. S., Chen, J. T., Sabapathy, K., & Ruan, D. Y. (2007). JNK1 contributes to metabotropic glutamate receptor-dependent long-term depression and short-term synaptic plasticity in the mice area hippocampal CA1. European Journal of Neuroscience, 25(2), 391–396. 10.1111/J.1460-9568.2006.05300.X

Luo, Z., Li, K., Yu, Y., Liu, Y., Weng, H., Zeng, X. T., Zheng, Y., & Li, W. (2026). LDHB Deficiency in Fibroblasts Induces Lactate-Mediated Inflammatory Reprogramming That Promotes Breast Cancer Metastasis. Cancer Research, 86(11), 2703–2721. 10.1158/0008-5472.CAN-25-2792/774529/AM/LDHB-DEFICIENCY-IN-FIBROBLASTS-INDUCES-LACTATE

Mathiesen, S. N., Lock, J. L., Schoderboeck, L., Abraham, W. C., & Hughes, S. M. (2020). CNS Transduction Benefits of AAV-PHP.eB over AAV9 Are Dependent on Administration Route and Mouse Strain. Molecular Therapy. Methods & Clinical Development, 19, 447. 10.1016/J.OMTM.2020.10.011

Meier, F., Brunner, A. D., Frank, M., Ha, A., Bludau, I., Voytik, E., Kaspar-Schoenefeld, S., Lubeck, M., Raether, O., Bache, N., Aebersold, R., Collins, B. C., Röst, H. L., & Mann, M. (2020). diaPASEF: parallel accumulation-serial fragmentation combined with data-independent acquisition. Nature Methods, 17(12), 1229–1236. 10.1038/S41592-020-00998-0

Mielke, K., & Herdegen, T. (2000). JNK and p38 stresskinases — degenerative effectors of signal-transduction-cascades in the nervous system. Progress in Neurobiology, 61(1), 45–60. 10.1016/S0301-0082(99)00042-8

Mohammad, H., Marchisella, F., Ortega-Martinez, S., Hollos, P., Eerola, K., Komulainen, E., Kulesskaya, N., Freemantle, E., Fagerholm, V., Savontous, E., Rauvala, H., Peterson, B. D., Van Praag, H., & Coffey, E. T. (2018). JNK1 controls adult hippocampal neurogenesis and imposes cell-autonomous control of anxiety behaviour from the neurogenic niche. Molecular Psychiatry, 23(2), 362–374. 10.1038/mp.2016.203

Oliva, A. A., Atkins, C. M., Copenagle, L., & Banker, G. A. (2006). Activated c-Jun N-Terminal Kinase Is Required for Axon Formation. The Journal of Neuroscience, 26(37), 9462. 10.1523/JNEUROSCI.2625-06.2006

Revest, J. M., Dupret, D., Koehl, M., Funk-Reiter, C., Grosjean, N., Piazza, P. V., & Abrous, D. N. (2009). Adult hippocampal neurogenesis is involved in anxiety-related behaviors. Molecular Psychiatry, 14(10), 959–967. 10.1038/MP.2009.15

Rincon, M. Y., De Vin, F., Duqué, S. I., Fripont, S., Castaldo, S. A., Bouhuijzen-Wenger, J., & Holt, M. G. (2018). Widespread transduction of astrocytes and neurons in the mouse central nervous system after systemic delivery of a self-complementary AAV-PHP.B vector. Gene Therapy 2018 25:2, 25(2), 83–92. 10.1038/s41434-018-0005-z

Sabio, G., Cavanagh-Kyros, J., Barrett, T., Jung, D. Y., Ko, H. J., Ong, H., Morel, C., Mora, A., Reilly, J., Kim, J. K., & Davis, R. J. (2010). Role of the hypothalamic–pituitary– thyroid axis in metabolic regulation by JNK1. Genes & Development, 24(3), 256. 10.1101/GAD.1878510

Samulski, R. J., & Muzyczka, N. (2014). AAV-Mediated Gene Therapy for Research and Therapeutic Purposes. 10.1146/annurev-virology-031413-085355

Scharfman, H. E. (2016). The enigmatic mossy cell of the dentate gyrus. Nature Reviews. Neuroscience, 17(9), 562. 10.1038/NRN.2016.87

Scharfman, H. E., & Myers, C. E. (2013). Hilar mossy cells of the dentate gyrus: a historical perspective. Frontiers in Neural Circuits, 6(DEC). 10.3389/FNCIR.2012.00106

Semba, T., Sammons, R., Wang, X., Xie, X., Dalby, K. N., & Ueno, N. T. (2020). JNK Signaling in Stem Cell Self-Renewal and Differentiation. International Journal of Molecular Sciences, 21(7), 2613. 10.3390/IJMS21072613

Sjöstedt, E., Zhong, W., Fagerberg, L., Karlsson, M., Mitsios, N., Adori, C., Limiszewska, A., Hikmet, F., Huang, J., Du, Y., Lin, L., Dong, Z., Yang, L., Liu, X., Jiang, H., Xu, X., Wang, J., Yang, H., Bolund, L., … Mulder, J. (2020a). An atlas of the protein-coding genes in the human, pig, and mouse brain. Science (New York, N.Y.), 367(6482). 10.1126/SCIENCE.AAY5947

Sjöstedt, E., Zhong, W., Fagerberg, L., Karlsson, M., Mitsios, N., Adori, C., Limiszewska, A., Hikmet, F., Huang, J., Du, Y., Lin, L., Dong, Z., Yang, L., Liu, X., Jiang, H., Xu, X., Wang, J., Yang, H., Bolund, L., … Mulder, J. (2020b). An atlas of the protein-coding genes in the human, pig, and mouse brain. Science (New York, N.Y.), 367(6482). 10.1126/SCIENCE.AAY5947

Tiwari, V. K., Stadler, M. B., Wirbelauer, C., Paro, R., Schübeler, D., & Beisel, C. (2011). A chromatin-modifying function of JNK during stem cell differentiation. Nature Genetics 2011 44:1, 44(1), 94–100. 10.1038/ng.1036

Tsuruta, F., Sunayama, J., Mori, Y., Hattori, S., Shimizu, S., Tsujimoto, Y., Yoshioka, K., Masuyama, N., & Gotoh, Y. (2004). JNK promotes Bax translocation to mitochondria through phosphorylation of 14-3-3 proteins. The EMBO Journal, 23(8), 1889. 10.1038/SJ.EMBOJ.7600194

Tyanova, S., Temu, T., Sinitcyn, P., Carlson, A., Hein, M. Y., Geiger, T., Mann, M., & Cox, J. (2016). The Perseus computational platform for comprehensive analysis of (prote)omics data. Nature Methods, 13(9), 731–740. 10.1038/NMETH.3901

Vicidomini, C., Guo, N., & Sahay, A. (2020). Communication, Cross Talk, and Signal Integration in the Adult Hippocampal Neurogenic Niche. Neuron, 105(2), 220–235. 10.1016/j.neuron.2019.11.029

von Ziegler, L. M., Floriou-Servou, A., Waag, R., Das Gupta, R. R., Sturman, O., Gapp, K., Maat, C. A., Kockmann, T., Lin, H. Y., Duss, S. N., Privitera, M., Hinte, L., von Meyenn, F., Zeilhofer, H. U., Germain, P. L., & Bohacek, J. (2022). Multiomic profiling of the acute stress response in the mouse hippocampus. Nature Communications, 13(1). 10.1038/S41467-022-29367-5

Winchester, C. L., Ohzeki, H., Vouyiouklis, D. A., Thompson, R., Penninger, J. M., Yamagami, K., Norrie, J. D., Hunter, R., Pratt, J. A., & Morris, B. J. (2012). Converging evidence that sequence variations in the novel candidate gene MAP2K7 (MKK7) are functionally associated with schizophrenia. Human Molecular Genetics, 21(22), 4910–4921. 10.1093/hmg/dds331

Wu, C. C., Tsantilas, K. A., Park, J., Plubell, D., Sanders, J. A., Naicker, P., Govender, I., Buthelezi, S., Stoychev, S., Jordaan, J., Merrihew, G., Huang, E., Parker, E. D., Riffle, M., Hoofnagle, A. N., Noble, W. S., Poston, K. L., Montine, T. J., & MacCoss, M. J. (2025). Enrichment of extracellular vesicles using Mag-Net for the analysis of the plasma proteome. Nature Communications, 16(1). 10.1038/S41467-025-60595-7

Yeh, C. Y., Asrican, B., Moss, J., Quintanilla, L. J., He, T., Mao, X., Cassé, F., Gebara, E., Bao, H., Lu, W., Toni, N., & Song, J. (2018). Mossy Cells Control Adult Neural Stem Cell Quiescence and Maintenance through a Dynamic Balance between Direct and Indirect Pathways. Neuron, 99(3), 493–510.e4. 10.1016/j.neuron.2018.07.010

Zeke, A., Misheva, M., Reményi, A., & Bogoyevitch, M. A. (2016). JNK Signaling: Regulation and Functions Based on Complex Protein-Protein Partnerships. Microbiology and Molecular Biology Reviews : MMBR, 80(3), 793. 10.1128/MMBR.00043-14

Zhou, X., Yi, W., Zhi, Y., Yu, J., Lu, D., Luo, Z., Yuan, L., Chen, L., Xu, Z., & Xu, D. (2023). Stress-Activated Protein Kinase JNK Modulates Depression-like Behaviors in Mice. Molecular Neurobiology, 60(5), 2367–2378. 10.1007/S12035-023-03209-X

Zincarelli, C., Soltys, S., Rengo, G., & Rabinowitz, J. E. (2008). Analysis of AAV Serotypes 1–9 Mediated Gene Expression and Tropism in Mice After Systemic Injection. Molecular Therapy, 16(6), 1073–1080. 10.1038/MT.2008.76

