## Supplementary figures for "Targeting distinct neuronal populations with AAV vectors reveals neurogenesis-associated anxiolytic effects of JNK inhibition"

**Supplementary figure 1**

**a)** Distance moved in the novel object recognition test of the mDlx-NLS-mRuby2/-JBD mice. **b)** Distance moved in the open field test of the mDlx-NLS-mRuby2 and mDlx-NLS-mRuby2-JBD mice at 10 weeks after AAV administration. Statistical analysis was done with Student's two-tailed t-test.

**Supplementary figure 2**

**a)** Timeline depicting the behaviour tests in the stress test experiments of AAV-PHP.eB-hSyn1/-NLS-JBD mice ("PHP.eB-hSyn1 groups"), AAV.CAP-B10-hSyn1-mRuby2/-NLS-JBD mice ("CAP-B10-hSyn1 groups"), and AAV.CAP-B10-mDlx-NLS-mRuby2/-JBD mice ("CAP-B10-mDlx groups"). PHP.eB-hSyn1 groups were tested at baseline and after stress exposure, after which the mice were sacrificed and blood was collected for analysis. CAP-B10-hSyn1 and CAP-B10-mDlx groups were tested at baseline, after stress, and the following day to measure recovery, after which the mice were sacrificed and blood was collected for analysis. **b)** Distance moved and **c)** time spent in the center in the open field test as well as **d)** spontaneous alternation in the Y-maze test were not affected by the stress exposure and expression of JNK inhibitor in the PHP.eB-hSyn1 mice. **e-g)** In CAP-B10-hSyn1 mice, there were no differences between the groups at baseline, after stress, or at recovery in **e)** distance moved and **f)** time spent in center in the open field test or **g)** in spontaneous alternation in the Y-maze test. **h-j)** In CAP-B10-mDlx mice, mice expressing JBD moved less in the open field at baseline **h)**, however, there were no significant differences after stress or at recovery. **i)** In these mice, there were no differences in time spent in the center of the open field or **j)** in spontaneous alternation in the Y-maze test at any of the three timepoints. Data show mean +/-SEM for n=12 PHP.eB-hSyn1-mRuby2 mice, n=14 PHP.eB-hSyn1-mRuby2-NLS-JBD mice, n=13 CAP-B10-hSyn1-mRuby2 mice, n=13 CAP-B10-hSyn1-mRuby2-NLS-JBD mice, n=13 CAP-B10-mDlx-NLS-mRuby2 mice, n=13 CAP-B10-mDlx-NLS-mRuby2-JBD mice. Statistical analysis was done with Student's two-tailed t-test. Abbreviations: FST=forced swim test, OF=open field, YM=Y-maze.

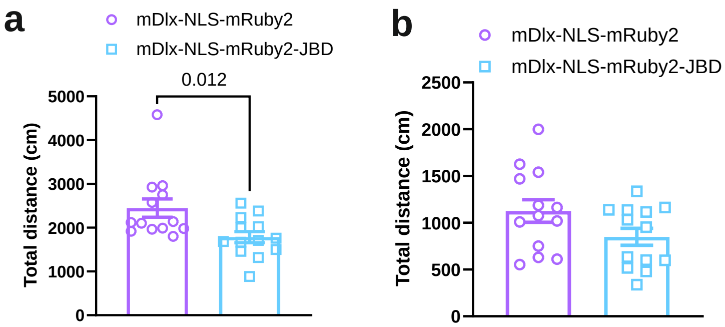


### Supplemental figure 1

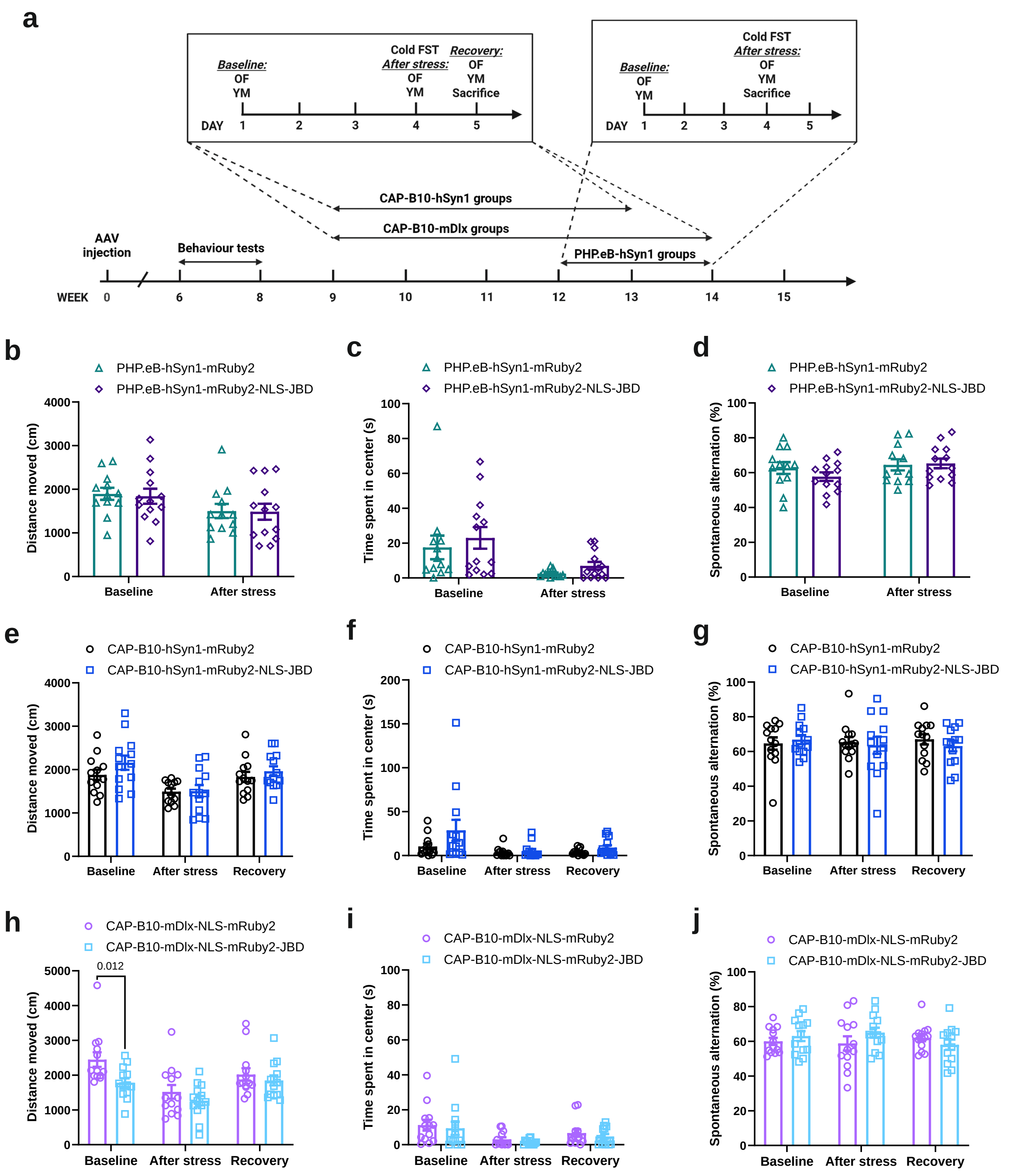


**Supplemental figure 2**
